# Unveiling Hidden Microbial Diversity in Mars 2020 Mission Assembly Cleanrooms with Molecular Insights into the Persistent and Perseverance of Novel Species Defying Metagenome Sequencing

**DOI:** 10.1101/2025.01.21.633868

**Authors:** Shobhan Karthick Muthamilselvi Sivabalan, Varsha Vijayakumar, Pratyay Sengupta, Siddhakam Palmal, Srinivasan Krishnamurthi, Nitin Kumar Singh, Nikos C. Kyrpides, Karthik Raman, Kasthuri Venkateswaran

## Abstract

NASA cleanrooms, where space mission components are assembled, maintain stringent cleaning protocols and nutrient-poor environments, resulting in low yet persistent microbial loads. Although these oligotrophic extremophiles are reported in small numbers, their resistance to environmental stresses, sparse presence, and difficulty in extracting biomolecules often lead to their omission, even with advanced sequencing technologies. Traditional metagenomic approaches fail to detect these rare species due to challenges in lysing robust microbial cells and isolating minute amounts of DNA from dominant microorganisms. Additionally, the absence of database references for novel extremophiles limits their identification. Over a six month period of monitoring Mars 2020 mission cleanrooms, 182 bacterial strains from 19 families were identified using advanced molecular techniques. This included 14 novel Gram-positive species, eight of which were spore-formers. Despite being present at only about 0.001% abundance in metagenomic sequencing data, they were successfully cultured. Functional studies revealed their capabilities in nitrogen cycling, carbohydrate metabolism, and radiation resistance. Furthermore, 12 biosynthetic gene clusters, including those linked to ectoine and *ɛ*-poly-L-lysine production, underscore their biotechnological potential. These findings emphasize the hidden microbial diversity in spacecraft assembly cleanrooms and highlight the need for advanced detection methods to uncover extremophiles with potential applications in biotechnology and space exploration.

**Synopsis:** Understanding extremophiles in NASA spacecraft assembly cleanrooms aids contaminant management in confined habitats, ensuring sustainability and safety in future space missions.

## Introduction

Many microorganisms, particularly those with specific growth needs or those that cannot survive outside their natural habitats, remain undetectable through standard culturing techniques. This is manifest in the difference between the microscope-aided microbial cell count and microbes that can be cultured using traditional laboratory methods, referred to as “Plate Count Anomaly”.^1^ Recent advancements in multi-omics analysis have shown the complicated and detailed aspects of microbial diversity found in specialized built environments, such as cleanrooms where pharmaceutical, medical, semiconductor, and aeronautical components are assembled. The contamination control strategies used to maintain these environments reduce microbial presence, paradoxically containing a diverse yet hidden array of microorganisms. This “rare/hidden microbiome” includes several low-abundance species that escape detection even by state-of-the-art molecular methods. These dormant forms, found in low-nutrient cleanrooms and present at low abundances, pose significant challenges for molecular detection using standard techniques.

Bacterial spores and *Actinobacteria* are robust, hard to lyse, and scarcely populated in oligotrophic environmental samples.^2^ NASA cleanrooms are low in biomass (∼100 viable cells or *<*1 cultivable cell per 25 cm^2^;^3^) and are considered an extreme oligotrophic environment due to the low number of total organic content (*<*1 pg per m^2^), maintaining cleanliness at ISO-6 level (*<*35,200 particles of 0.5 µm per feet^3^), and less availability of nutrients to microorganisms.^4^ Given stringent contamination control protocols, extremophiles in cleanrooms are less metabolically active; consequently, their genetic footprints are often underrepresented in metagenome sequencing data. Moreover, current DNA extraction methods are typically optimized for the most abundant or robust microbial populations, often overlooking rare microbes (*<*1%). This bias arises because standard protocols may not effectively lyse all cell types or capture DNA from low-abundant microorganisms, leading to underrepresentation of rare species in metagenomic analyzes.^5^ Further, insufficient sequencing depth during shotgun metagenomics exacerbates the issue, as the sequencing effort is disproportionately distributed among the dominant species.^6^ Biases introduced during DNA amplification and library preparation can also skew the results, failing to accurately reflect the true microbial diversity.^7^ To rectify these issues, it is essential to develop more inclusive DNA extraction protocols that ensure efficient DNA recovery from all types of microbial cells, increase the depth of sequencing, and employ techniques that minimize amplification biases.^8^ Another significant limitation of metagenomic sequencing is its reliance on comprehensive reference databases for species identification. The genetic sequences of certain extremophiles have hitherto not been cataloged and, hence, not widely present in reference databases. Therefore, they may continue to remain undetected, underlining the need for continual updates to these databases with genetic information from newly sequenced extremophiles. These improvements are crucial for comprehensively and accurately characterizing microbial communities, particularly in environments where rare microbes play significant ecological roles.

A key objective of this study was to sequence the genomes of the 182 isolated bacterial strains and to delineate their phylogenetic affiliations within the Mars 2020 mission assembly cleanrooms.^9–11^ To fully understand the taxonomic and evolutionary background of these novel species, a comprehensive phylogenomic investigation was carried out using multiple biomarkers, and a thorough evaluation of their physiological characteristics was performed. In conjunction with genomic characterization, the metagenomic signatures derived from the cleanroom environment were also investigated to determine the prevalence, incidence, and persistence of these novel species throughout the critical phase of the Mars 2020 mission assembly. A parallel line of inquiry involved probing into the functional content of these novel species, with a keen focus on discovering unique enzymes and secondary metabolites. Such metabolites could provide valuable clues regarding the adaptive mechanisms of these rare novel species, enabling them to survive in the nutrient-scarce conditions of the cleanroom.

Ultimately, by mapping these survival strategies and functional potentials, the study unearthed insights into how these hidden extremophilic bacteria manage to thrive within the unique conditions of the cleanroom. Figure 1 shows the overall workflow of this study, outlining the key analyses performed, from sample collection and microbial isolation to genome sequencing, phylogenetic classification, and functional characterization of novel bacterial species. Overall, this study enriches our understanding of microbial resilience and adaptability and bears significant implications for future space exploration.

**Figure 1:**
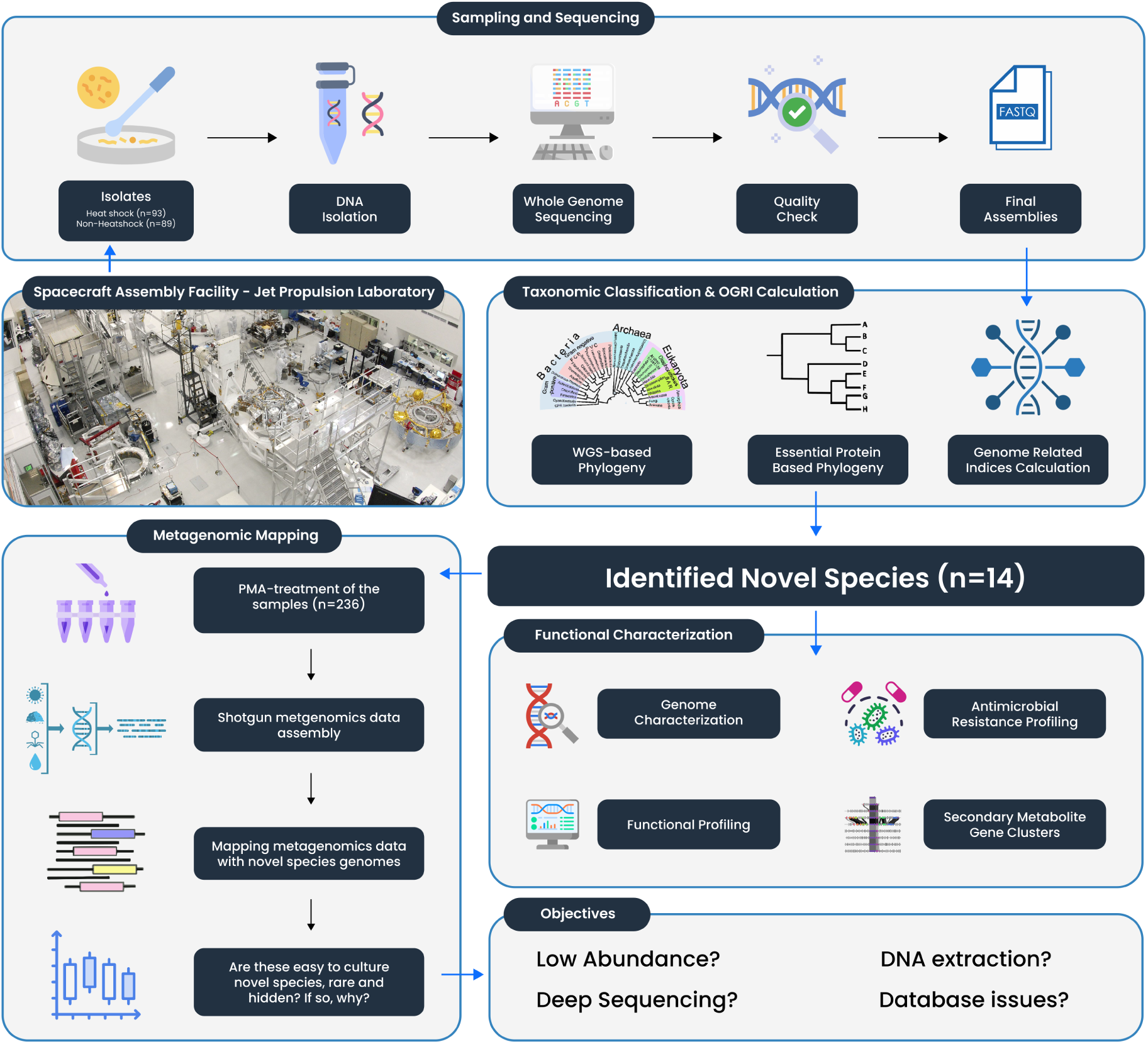
Overview of the study design including I. Sampling and sequencing procedures, II. Taxonomic classification, III. Mapping of metagenomic reads to estimate the abundance of novel species, and IV. Functional characterization. JPL-SAF Photo credit: NASA/JPL-Caltech

## Results

### Dominance of spore formers in Mars 2020 assembly cleanroom

During the microbial surveillance of the Mars 2020 assembly cleanroom at JPL-SAF, 182 bacterial strains were isolated (Supplemental File S1) and the whole genome sequencing (WGS) classification established them into 19 bacterial families (Table 1). Seasonal and temporal distribution analysis showed that these cleanroom bacterial strains were not abundant during the 6-month period or at any location, since no definitive pattern was observed (Supplemental File S2). Among 92 isolates obtained after the heat shock (HS) procedure (HS; 80*^◦^*C for 15 min), 17 genera and 32 species were identified. Heat shock procedure eliminated the majority of the non-spore-forming bacteria and 93.5% (n = 86 isolates) were spore-forming. It is interesting to emphasize that six non spore-forming strains belonging to the *Brevibacteriaceae*, *Micrococcaceae* and *Staphylococcaceae* families could withstand the HS procedure, and their heat tolerance was further confirmed by exposing them again at 80*^◦^*C for 15 min. Approximately 66% of the strains from the HS samples belong to the *Bacillaceae* family (n = 61 isolates), comprising 11 genera and 23 species. The second dominant family was *Brevibacillaceae* (n = 12 isolates), with one genus and three species. At the species level, *Bacillus pumilus* and *Bacillus subtilis* were dominant in HS samples. Further molecular analyzes revealed the presence of seven novel species; four of them were members of *Bacillaceae*; two from *Planococcoceae*, and one from *Brevibacillaceae*, which are described in this study using molecular phylogeny.

**Table 1.**
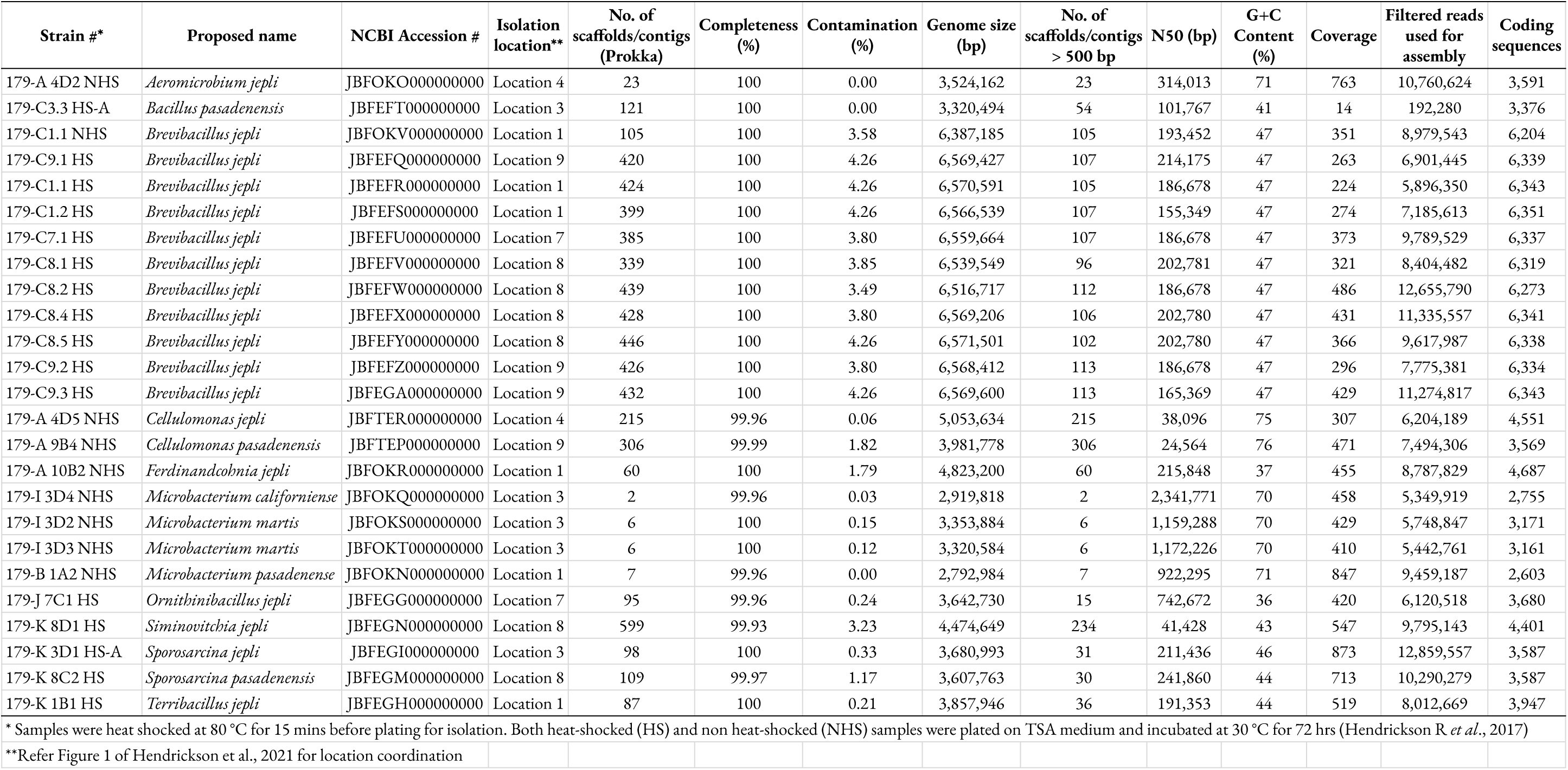
Genome assembly and annotation metrics of the novel species described during this study.

Unlike the HS bacterial isolates, the 90 strains recovered from non-heat-shocked (NHS) samples revealed a diverse range of taxa, covering 30 genera and 57 species. Approximately 52% of the NHS strains were not producing spores. However, except for five strains that belong to Gram-stain negative members of *Sphingomonadaceae*, all other NHS isolates were Gram-stain positive. Strains belonging to the *Bacillaceace* family (n = 28 isolates) were dominant in NHS samples that consisted of four genera and 14 species. The second dominant members belong to the *Micrococcaceae* family (n = 16 isolates), human skin microbes. At the species level, *Bacillus safensis* (n = 6 isolates) and *Bacillus altitudinis* (n = 5 isolates) were dominant in NHS samples. Further molecular analysis revealed the presence of seven novel species after NHS treatment. One novel species each *Bacillaceae* and *Nocardioodaceae*; two from *Cellulomonadeceae*; and three from *Microbacteriaceae* were identified. One novel species of the genus *Brevibacillus* was retrieved from both NHS and HS samples. In total, 14 novel bacterial species are described, all of which are Gram-stain positive. Among these, eight species are forming spores while the remaining six species are unable to produce spores.

### Genome relatedness indices confirmed 14 novel bacterial species

The assembly statistics of the novel species, such as genome size, number of contigs, and the N50 metric, along with metadata, are provided in Table 1, obtained through the quality assessment of the assembled genomes using QUAST v.5.0.2. Evaluation of genome assemblies for completeness and contamination reveals that genomes of most strains exhibited completeness *>*99% and contamination *<*4.26%. Based on marker genes (16S rRNA, gyrB), the similarities between the novel species and closely related NCBI species, the average amino acid identity (AAI), the average nucleotide identity (ANI) and the digital DNA-DNA hybridization (dDDH) are presented in Table 2. As established, ANI values less than 95% and a dDDH value below 70% are strict thresholds for the definition of new species.^12^ The results in Table 2 for the species follow these criteria, confirming that they are novel. The ANI values for WGS and the closest validly described species range from *<*77% to 94.12%. Furthermore, for WGS, the dDDH indices ranged from 19. 5% to54. 2%. Since there is no universally established threshold for the classification of bacterial genus levels based on AAI, gyrB, and 16S rRNA similarity, it is challenging to definitively determine whether these species belong to novel genera.

**Table 2.**
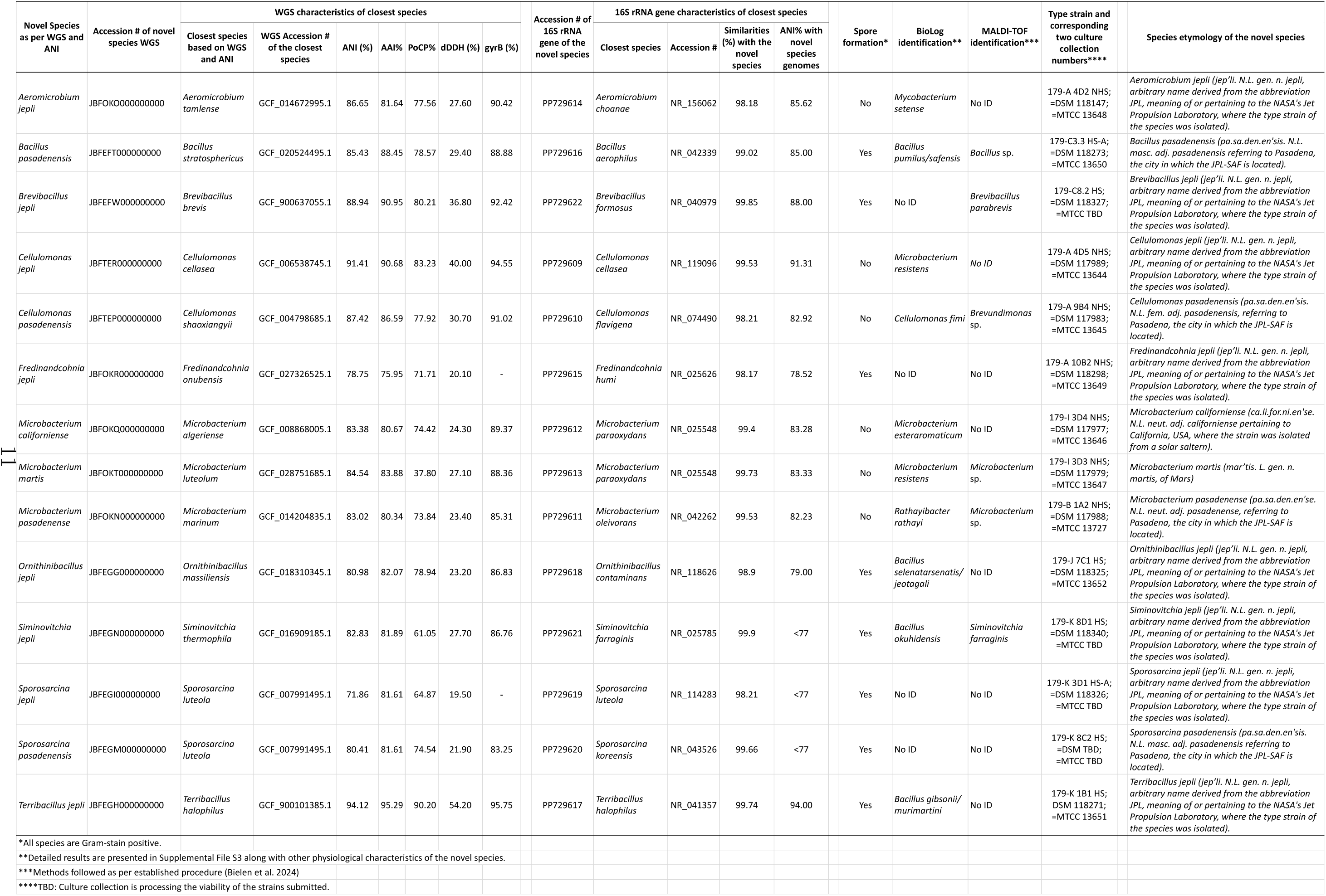
Taxonomic identification, similarities of various gene markers, overall genome-relatedness indices, dDDH analysis, and etymology of novel bacterial species.

### Phylogenomic analysis revealed diversity of novel species in cleanroom isolates

Individual phylogenetic trees were constructed for each novel genome at the genus level to understand their evolutionary history and relatedness to already recognized genomes and newly identified species. The taxonomic placement of each novel genome was further compared with the closest species identified by WGS by ANI. To confirm the placement of the novel species within the bacterial tree of life, a comprehensive phylogenetic tree was generated using 5,489 complete, representative, non-anomalous bacterial genomes along with the 14 novel species (n = 25 strains). The diversity of bacteria present in cleanrooms is clearly reflected in the tree, as newly identified species are distributed across multiple branches of the bacterial tree of life. Individual trees for each novel species, along with the bacterial tree of life, are shown in Figures 2A to 2J.

**Figure 2:**
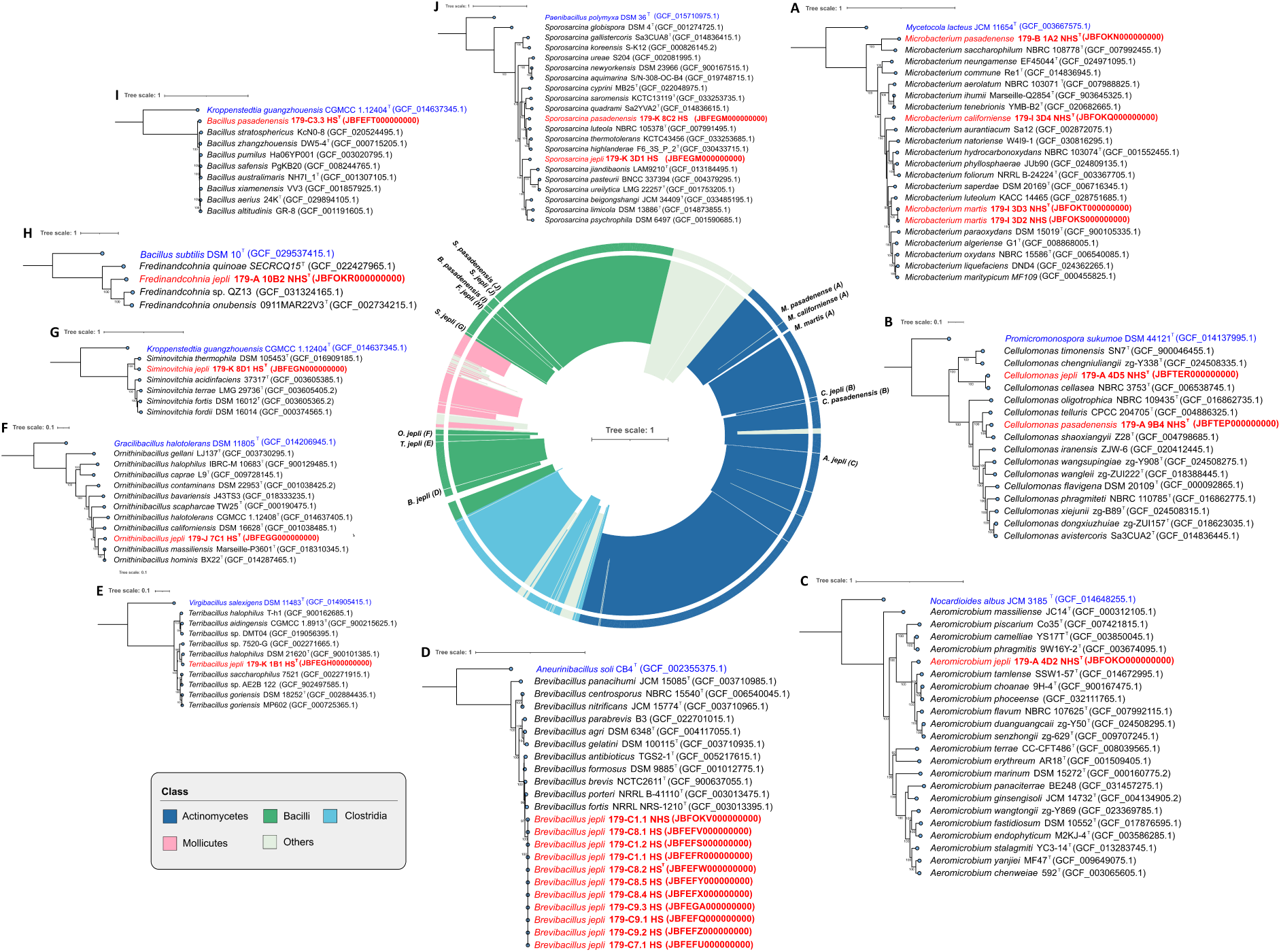
The phylogenetic placement of novel bacterial taxa identified in this study. Phylogenetic trees (A-J) were constructed using a set of available marker sequences. The central tree of life indicates the position of novel species in the bacterial tree of life. Novel species and the outgroups are highlighted in red and blue, respectively, and their corresponding NCBI accessions are provided.

Six novel species were recognized under the *Actinomycetota* phylum (Figure 2A to 2C). Among the four strains belonging to the genus *Microbacterium*, three were identified as novel species (Figure 2A). The two strains of *Microbacterium martis* shared 100% similarity with each other and exhibited ANI values of 78% to 85% with validly established species. For *Microbacterium pasadenense* 179-B 1A2 NHS, the nearest species was *Microbacterium dauci* with 83.21% ANI similarity, while *Microbacterium saccharophilum* was identified as the closest species based on single-copy genes. *Microbacterium californiens*e 179-I 3D4 NHS showed ANI values between 85% and 78% with representative genomes from NCBI, with *Microbacterium galbinum* being the closest species. For *Cellulomonas jepli* 179-A 4D5 NHS, the closest species based on ANI was *Cellulomonas cellasea* with 91.43%, while for *Cellulomonas pasadenensis* 179-A 9B4 NHS, it was *Cellulomonas shaoxiangyii* with 87.52% ANI (Figure 2B). *Aeromicrobium jepli* 179-A 4D2 NHS exhibited ANI values ranging between 87% and 79% with recognized *Aeromicrobium* species, with *Aeromicrobium tamlense* being the closest at 86.73% similarity (Figure 2C).

Members of the *Firmicutes* phylum exhibited a wide range of ANI values with recognized species (n = 8). The 11 strains of *Brevibacillus jepli* clustered together and showed *Brevibacillus brevis* type strain as the nearest species with 89.03% ANI (Figure 2D), though analysis based on single-copy genes indicated *Brevibacillus fortis* as the closest relative. *Terribacillus jepli* 179-K 1B1 HS exhibited 94.17% ANI similarity with *Terribacillus halophilus* and ANI values ranging from 94% to 80% with other members of the same genus (Figure 2E). For *Ornithinibacillus jepli* 179-J 7C1 HS, the closest relative was *Ornithinibacillus hominis* with 80.91% ANI, and it also shared 80–77% similarity with other recognized species, clustering closely in the phylogenetic tree (Figure 2F). *Siminovitchia jepli* 179-K 8D1 HS had *Siminovitchia thermophila* as its closest relative with 82.69% ANI (Figure 2G). *Fredinandcohnia jepli* 179-A 10B2 NHS exhibited 78.76% ANI similarity with *Fredinandcohnia onubensis* (Figure 2H).

The novel species *Bacillus pasadenensis* 179-C3.3 HS-A was placed near *Bacillus zhangzhouensis* in the phylogenetic tree with 85.38% similarity, sharing ANI values ranging from 85% to 77% with other established species of the same genus (Figure 2I). Finally, the two novel species, *Sporosarcina jepli* and *Sporosarcina pasadenensis*, belonging to the genus *Sporosarcina*, exhibited ANI values of less than 77%, with their nearest valid species, *Sporosarcina luteola* and *Sporosarcina koreensis*, respectively (Figure 2J).

In addition to the phylogenomics analyses to identify the bacterial taxa, all novel species were subjected to BioLog phenotypic array^13^ and matrix-assisted laser desorption/ionization (MALDI) profile^14^ based bacterial identification and results are presented in Supplemental File S3. The type strains of all these novel species were deposited in two internationally recognized culture collections and details are provided in Table 2.

### Universal marker gene highlights rarity of novel species in existing databases

In this study, we collected the genes corresponding to the COG0013 (Alanyl-tRNA Synthetase, AlaS), a universal marker gene sequence,^15^ from 14 novel species using CD-HIT clustering to assess their similarity with WGS in the GTDB. The clustering of CD-HIT at an identity threshold 98% indicated that four of the 14 novel species described in this study had high sequence similarity to existing entries in GTDB. Three species that had higher CD-HIT values were *Sporosarcina* sp. Marseille-Q4943 (NCBI), *Cellulomonas* sp. NS3 (IMG, NCBI, metaG), and *Brevibacillus brevis* NRRL NRS-604 (IMG, NCBI, metaG). The fourth novel species with higher CD-HIT values exhibited *>*98% COG sequence similarities with *Aeromicrobium* sp. MAG from composted filter cake microbial communities from a cane sugar milling plant in Nova Europa, Brazil. To validate the CD-HIT findings, we performed pairwise ANI analyses between the WGS of the higher CD-HIT valued species and the corresponding novel strains. Pair-wise ANI analysis showed that *S. pasadenensis* 179-K 8C2 HS strain had 93% ANI with *Sporosarcina* sp. Marseille-Q4943; *C. jepli* 179-A 4D5 NHS had only 93% ANI with *Cellulomonas* sp. NS3 (IMG,NCBI); *B. jepli* 179-C8.2 HS exhibited 89% with the type strain of *B. brevis* NRRL NRS-604; and ANI was *<*85% between *A. jepli* 179-A 4D2 NHS and *Aeromicrobium* sp. MAG, instituting the importance of a culture-based approach in studying previously predicted organisms. These ANI values are below the 95% threshold commonly used to delineate bacterial species,^16^ indicating that the 14 novel species identified in this study are distinct from previously described species and not found in any of the databases comprising 28,856 validly recognized species, or 88,437 MAGs, or 80,141 high-quality curated IMG-metagenomes (Figure 3A). The observed discrepancy in the CD-HIT analysis, where the novel strain 179-C8.2 HS exhibited high sequence similarity with *B. brevis* type strain NRRL NRS-604 based on AlaS protein sequences, necessitates further scrutiny when employing the CD-HIT approach. This underscores the importance of complementing protein sequence clustering with whole-genome analyses to accurately determine species novelty.

**Figure 3:**
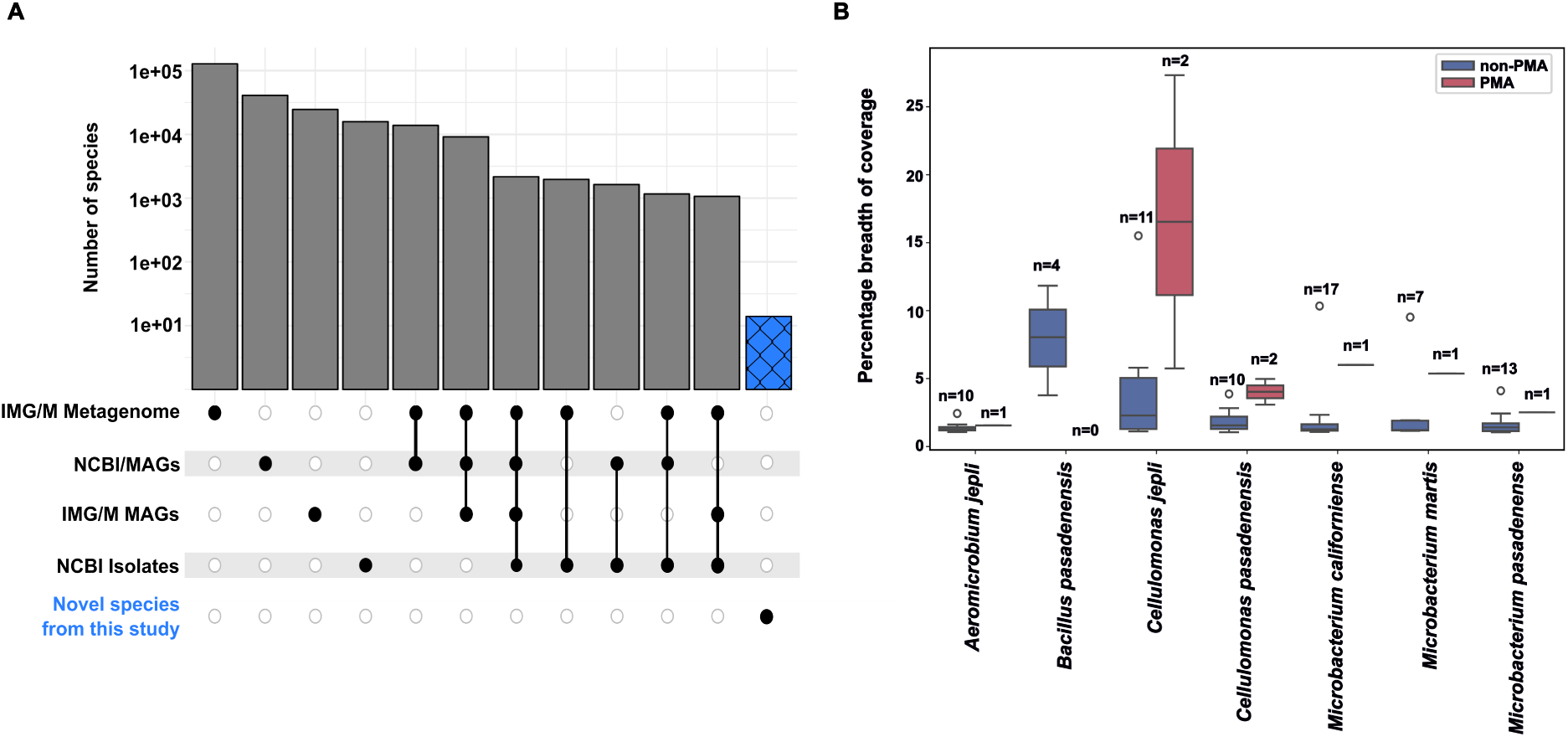
(A) Comparison of species detection across datasets. Bar plots display the number of species identified in various data sources: IMG/M metagenomes, NCBI/MAGs, IMG/M MAGs, and NCBI isolates. Blue bar represent novel species identified in this study. The Upset plot beneath the bar chart indicates overlaps among data sources, with filled circles marking shared species between datasets. (B) Percentage breadth of genome coverage of identified species, compared between PMA-treated (pink) and non-PMA-treated (blue) samples. Box plots represent median coverage with whiskers indicating data variability. Specific species are labeled with their incidence (n).

### Novel species exhibits low abundance in cleanroom metagenome

Shotgun metagenomic reads (n = 236 samples) from different locations within JPL-SAF were mapped to the novel species genome to determine their abundance and prevalence within the cleanroom environment. The evaluation criteria were *>* 10,000 bp as aligned assembled length, *>*100 mapped reads, and *>*1% breadth of coverage of the genome. The distribution of the breadth of coverage for novel species that passed filtering for at least one sample is shown in Figure 3B. A comparison between propidium monoazide (PMA) and non-PMA samples revealed significantly fewer PMA samples, all of which were assigned to Location 9 in JPL-SAF. The average breadth of coverage ranged between 1.05% and 15.51% for non-PMA samples and 1.55% to 27.32% for PMA samples. For the new species, the highest breadth of coverage was in *C. jepli* for the PMA and non-PMA samples, *M. californiense* and *M. martis* were next for the PMA samples, and *B. pasadenensis* and *M. californiense* followed for the non-PMA samples. Seven novel species (*B. jepli*, *T. jepli*, *O. jepli*, *Sim. jepli*, *Fredinandcohnia jepli*, *Spo. pasadenensis* and *Spo. jepli*) had a breadth of coverage *<*1% and were therefore excluded from the figure. These results of the metagenome mapping indicate that the abundance of the novel species within JPL-SAF is extremely low.

### Novel species exhibit diverse stress, biofilm, and metabolic adaptations

The findings of the functional characterization of the 14 novel species are summarized in Figure 4, which involved annotating their genomes using Prokka and classifying them using cogclassifier and Distilled and Refined Annotation of Metabolism (DRAM). Cellular processes, environmental information processing, genetic information processing, metabolic, and unknown were the five primary functional categories into which the COGs from the genomes were categorized (Supplemental File S4). Many genes were classified for amino acid transport and metabolism (389), transcription (344), and carbohydrate transport and metabolism (274).

**Figure 4:**
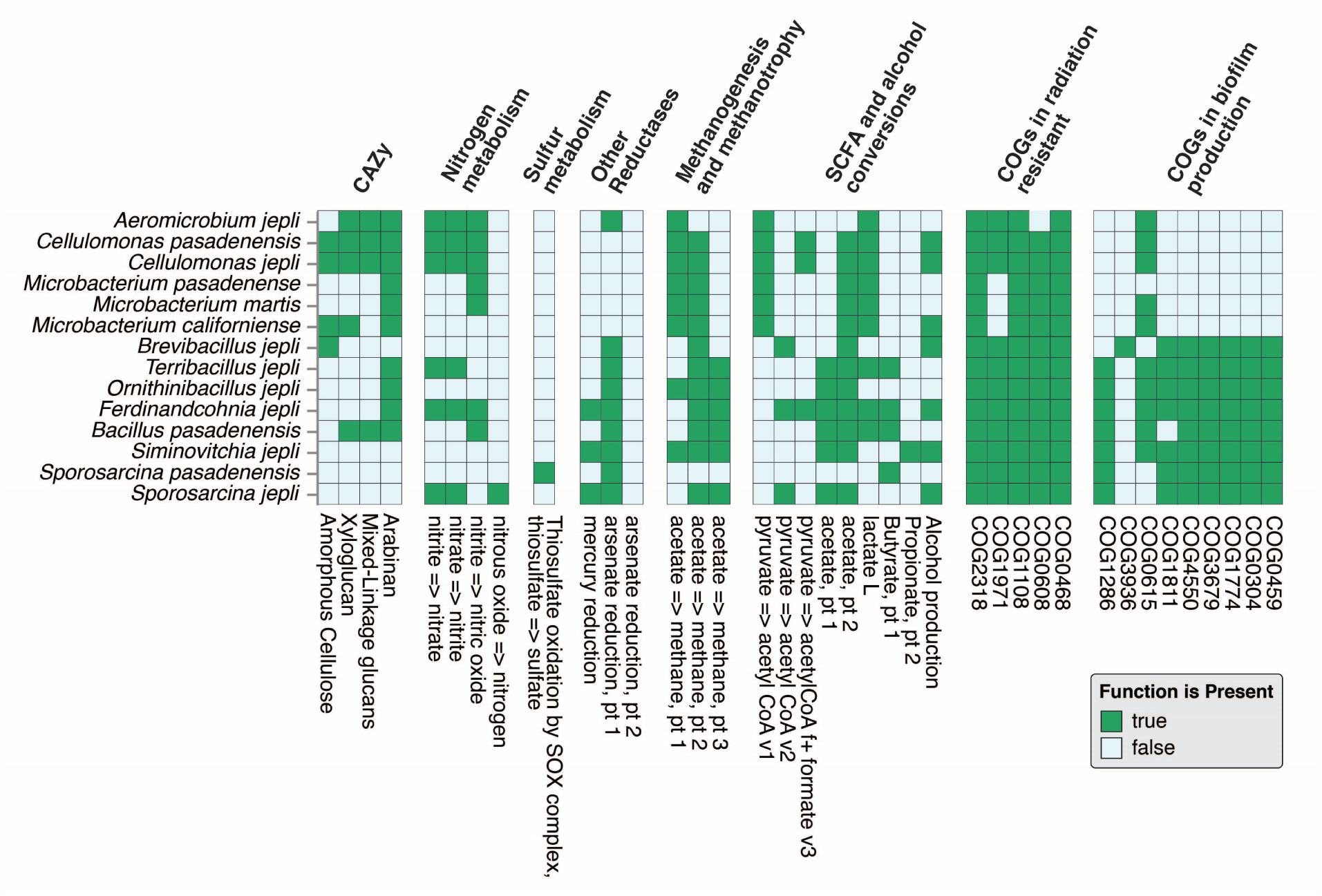
Functional profiling of identified species. Heatmap illustrates the presence or absence of metabolic pathways and functional genes across bacterial species. Functional categories include carbohydrate-active enzymes (CAZy), nitrogen metabolism, sulfur metabolism, reductases, methanogenesis and methanotrophy, SCFA (short-chain fatty acid) and alcohol conversions, radiation resistance (COGs), and biofilm production (COGs). The y-axis lists species (ordered phylogenetically), while the x-axis represents the functional categories and specific pathways.

The stress tolerance mechanisms in these novel genomes can be attributed to the presence of radiation resistant genes. The homologous recombination process by which DNA damage is repaired included key genes in all species, namely COG0468 (RecA) and COG2318 (DinB). Zinc uptake systems consisting of membrane-bound proteins and transcription regulation under stress conditions were found to occur in all species with COG1108 (ZnuB). However, COG0608 - RecJ, involved in DNA recombination and repair, was not present in *A. jepli*. Likewise, proteins such as DNA repair proteins, COG1971, or MntP, were absent in nonspore-forming species of the *Microbacterium* genus. Such observations highlight the diversity of the stress tolerance and repair mechanisms associated with the new genomes. The genes involved in biofilm production were found mainly in novel species that form spores. COG0459 (GroEL), a chaperonin protein, and COG0304 (FabB), which is involved in fatty acid biosynthesis and is essential for the formation of biofilm structure, were found in all spore formers but not in non-spore formers. Similarly, the tripartite complex YaaT/YliF/YmcA (COG1774, COG3670, and COG3679), which enhances biofilm formation by promoting Spo0A (a transcriptional regulator), was found exclusively in spore-forming species. However, neither *B. pasadenensis* nor any non-spore formers exhibited COG1811 (YqgA), a gene that affects biofilm formation. The contributor to biofilm production COG0615, also known as TagD, was lacking in both *Sporosarcina* species, *B. jepli* and *M. pasadenensis*. The accessory protein, CvpA (COG1286), which contributes to colicin V production and biofilm formation, was present in all spore formers except *B. jepli*. Finally, COG3936, an acyltransferase with localization inside the membrane, was only present in *B. jepli*. These results underscore the important role of spore-forming mechanisms in the facilitation of biofilm production among these novel species.

Looking at the results of DRAM, an analysis of carbohydrate-active enzymes (CAZy) for the novel species revealed that it possesses a wide range of metabolic capabilities, mainly for the degradation of polysaccharides, as illustrated in Figure 4. In these, genes for arabinan degradation were present in almost all species. It was observed that in *C. pasadenensis* and *C. jepli*, there is an increased repertoire of genes that allow the breakdown of arabinan, xyloglucan, amorphous cellulose, and mixed-linkage glucans. In nitrogen metabolism, genes related to the conversion of nitrite to nitric oxide, which is subsequently converted to nitrous oxide, and nitrate reduction pathways were identified within the genomes for nitrogen metabolism. Sulfur metabolism was indicated by the genes responsible for the oxidation of thiosulfate through the Sox complex, found exclusively in *Spo. pasadenensis*. In addition, metabolic profiles include genes related to short-chain fatty acids (SCFA), methanogenesis, and other reductases. The pathways for synthesis and conversion of acetate, propionate, and butyrate show that these novel species contribute to carbon flux in an anaerobic or microaerophilic environment. These findings suggest that these new species have a wide range of functional abilities in terms of environmental adaptation and nutrient cycling.

### Genomic insights into antibiotic resistance mechanisms in novel bacterial species

AMR analysis identified various gene families across the genomes, indicating resistance to nine different drug classes (Supplemental File S5). As depicted in Figure 5A, the highest prevalence of resistance to glycopeptides, fluoroquinolones, disinfecting agents, antiseptics and phenicol antibiotics was observed. Among the 14 novel species, *B. jepli* 179-C8.2 HS exhibited the highest number of AMR genes, followed by *Spo. pasadenensis* 179-K 8C2 HS, *O. jepli* 179-J 7C1 HS, *F. jepli* 179-A 10B2 NHS, and *Spo. jepli* 179-K 3D1 HS, all of which are spore-forming species. These AMR mechanisms could be classified into three types; the predominant type was alteration of antibiotic targets (Figure 5B). Other, less frequently found mechanisms were antibiotic efflux and antibiotic inactivation. The overall genomic analysis shows that there were 72 AMR genes distributed in all 14 genomes, though more phenotypic studies are needed to validate functionality.

**Figure 5:**
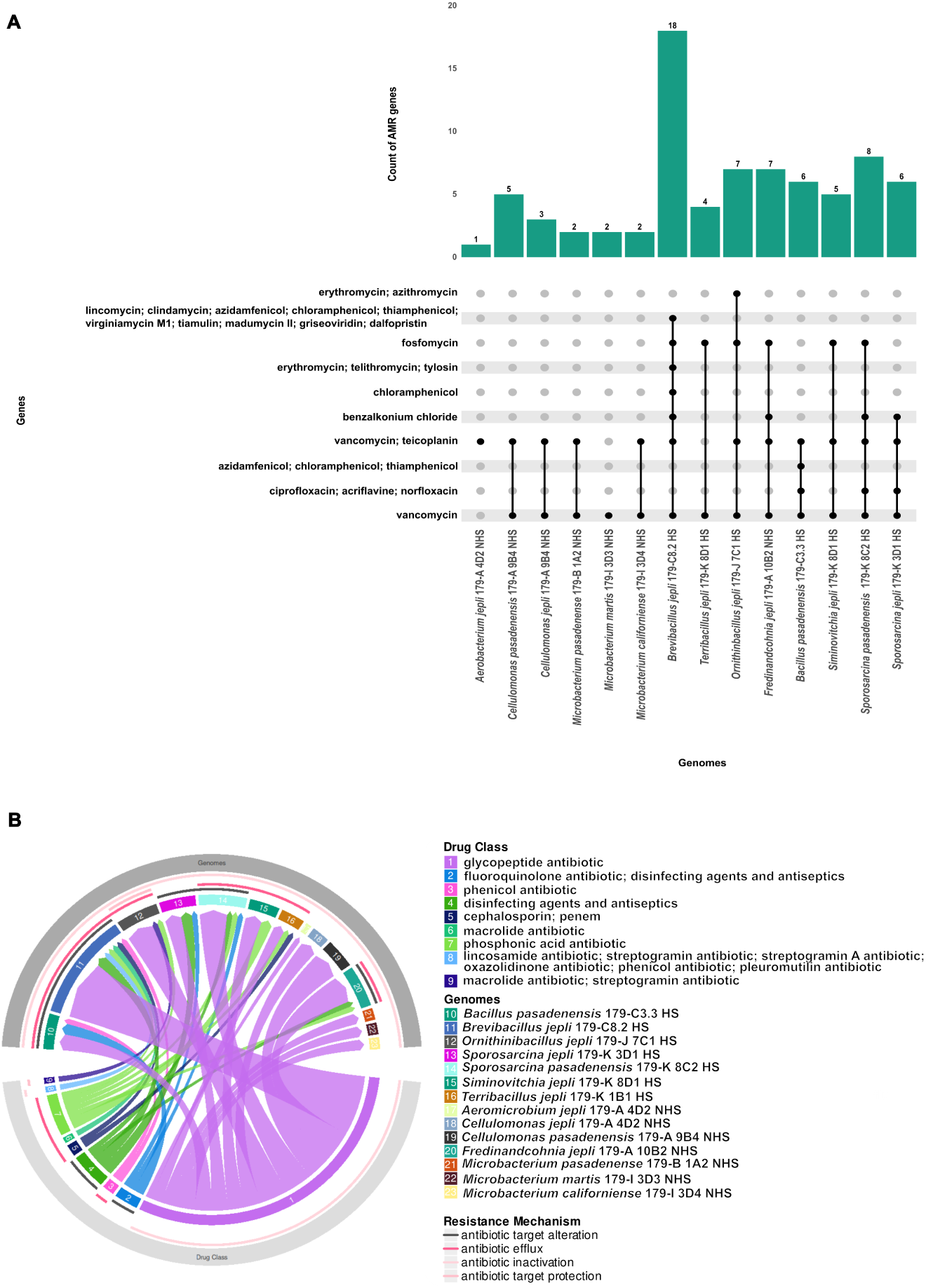
(A) Bar chart showing the number of antibiotic resistance genes (ARGs) detected in each of the 14 bacterial species isolated from the spacecraft assembly facility. Each bar represents the total number of ARGs found in the corresponding novel species. The upset plot shows the overlap of different antibiotic classes to which the ARGs provide resistance across the novel species. (B) Chord diagram illustrating the distribution of ARGs and their associated resistance mechanisms across the 14 bacterial species. The outer ring shows the different drug classes represented by ARGs, while the inner ring shows the bacterial genomes. Lines connecting the outer and inner rings indicate the presence of ARGs conferring resistance to specific drug classes in each species. The colors of the lines represent the different resistance mechanisms (e.g., antibiotic target modification, antibiotic inactivation).

### Biosynthetic gene clusters suggest biotechnological relevance of novel species

Analysis of biosynthetic gene clusters (BGC) showed that the 14 novel species had 12 different types of clusters, each of which was more than 60% related to existing BGCs (Supplemental File S6). The two most common clusters were *ɛ*-Poly-L-lysine (*ɛ*-PL) and ectoine which were *B. jepli* 179-C8.2 HS and *C. pasadenensis* 179-A 9B4 NHS which also showed the highest number of BGC. Among the identified clusters, six had more than 80% similarity to known BGC: ectoine, *ɛ*-Poly-L-lysine, petrobactin, ulbactin F / ulbactin G, bacilysin and alkylresorcinol. In particular, certain BGC isolates from *Spo. jepli*, *C. jepli*, *C. pasadenensis* and *B. jepli* showed 100% similarity to the ectoine, *ɛ*-poly-L-lysine, ulbactin F / ulbactin G and alkylresorcinol groups, respectively, thus highlighting the metabolic potential of these novel species for producing bioactive compounds.

Further detailed analysis revealed that both *Cellulomonas* species contain *ɛ*-PL (Figure 6). The *ɛ*-PL was also detected in several new non spore-forming species isolated from NASA clean rooms, which is 100% similar to the cluster of a fungus *Epichloe festucae* known for producing the natural homopolymer with antimicrobial properties used to preserve food, as well as for use in pharmaceutical applications. The presence of *ɛ*-PL BGC mediated through specific gene clusters underscores the potential of these novel species in terms of *ɛ*-PL production. Their high similarity to those of the already characterized clusters emphasizes their capability of producing bioactive compounds with significant applications in biotechnology and medicine. More research is needed with respect to the expression, regulation, and functional roles of these clusters to explore their entire biotechnological potential.

**Figure 6:**
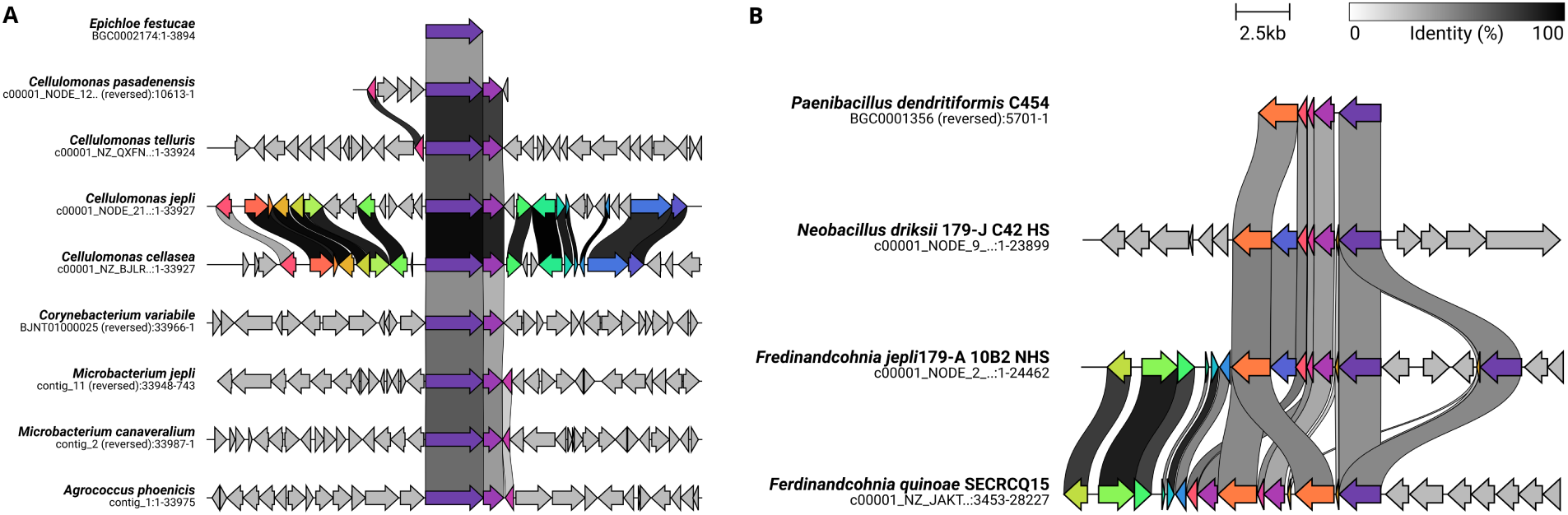
Comparative analysis of specialized metabolite biosynthetic gene clusters in novel species. (A) Conservation of the *ɛ*-Poly-L-lysine biosynthetic gene cluster across various species. The alignment compares the *ɛ*-Poly-L-lysine gene clusters from the known producers: *Epichloe festucae* (fungi), *Corynebacterium variabile* (bacteria), and the novel species identified from several NASA assembly facilities during this work (*Cellulomonas jepli*, *C. pasadenensis*) and in our previous studies (*Microbacterium jepli*, *M. canaveralium*, *Agrococcus phoenicis* ^17^), and their closest phylogenetic neighbors. Homologous regions are shown with shaded connectors, and arrows represent genes, color-coded by function, demonstrating a conserved gene cluster architecture among these species. (B) Comparative analysis of the paeninodin biosynthetic gene cluster. The alignment highlights conserved regions of the paeninodin cluster between *Paenibacillus dendritiformis* (a known producer) and the novel species (*Ferdinandcohnia jepli* and its closest neighbor*Ferdinandcohnia quinoae*). Shaded regions indicate sequence identity (%), with arrows showing the direction and function of genes in the cluster, revealing both conserved and unique genetic elements.

## Discussion

Unexplored microbiomes represent an untapped frontier of exploratory microbiology. As we delve into data-driven microbiome analysis, researchers are beginning to uncover microbial species that were previously invisible. The discovery of the 14 novel species and the elucidation of their metabolic pathways in one of the cleanest and most controlled anthropogenic systems was made possible only due to significant advancements in sample processing and data analysis strategies. Despite rapid progress in DNA isolation, culturomics, contamination control, and database-driven species identification, all these fields need further optimization to be leveraged as a strategic pipeline for pathogen microbiology, astro-microbiology, and epidemiology, where hidden microbes might decide the fate of a space mission or the health of a patient.

Extremophiles that are resilient and adapt to difficult-to-thrive environmental conditions defy conventional DNA extraction and sequencing methods. The cellular structures of these extremophiles are complex to lyse open and access to the biomolecules of interest, leading to underrepresentation in environmental samples and biases in molecular analyzes.^18,19^ Traditional laboratory techniques, such as the physicochemical disruption of these robust cells by boiling and freeze-thaw cycles, prove inadequate for the disruption of these robust cells.^20^ Such methods often fail to accurately represent less abundant species, leading to a skewed representation of the overall composition of the community. This bias can result in an underrepresentation of rare taxa, which ultimately affects the interpretation of microbial diversity and ecosystem dynamics.^21^ Deeper sequence coverage and rigorous bioinformatics might mitigate these biases. To understand functional predictions, it is important to update microbial reference databases with genomic data from novel extremophiles. Hence, it is critical to focus on developing novel nucleic acid extraction strategies by including enzymatic degradation of rigid cell walls of extremophiles (e.g. MetaPolyzyme ^22^), conducting deep sequencing, and improving reference databases by updating genomic information with novel extremophiles.

Culturing extremophiles in laboratory settings is essential for resistance testing, helping to develop effective cleaning and sterilization protocols.^3,23^ This study identified 14 novel bacterial species from the Mars 2020 assembly cleanrooms (∼14% of the 182 strains characterized), confirmed as distinct through ANI analysis, despite the high similarity of the AlaS protein sequence with recognized species (Figure 3A). Pairwise ANI values below the 95% threshold underscore the superiority of WGS over few protein-based clustering approaches for accurate taxonomy. Metagenomic mapping revealed an extremely low prevalence of these species in Mars 2020 assembly facilities, consistent with challenges in detecting rare microbes due to sequencing biases and limited reference databases (Segata et al., 2013). These results prove the rarity and transient nature of these novel species in cleanrooms, their adaptations to oligotrophic conditions, and the absence of clear temporal or seasonal patterns (Supplemental File S1).

The genomic and functional characterization of 14 newly identified bacterial species in this study revealed diverse metabolic capabilities and stress response mechanisms, underscoring their adaptability to extreme or nutrient-limited lithotrophic environments. The presence of CAZymes capable of degrading complex polysaccharides such as arabinan, xylan, and amorphous cellulose suggests that NASA cleanroom bacteria can utilize a wide range of carbohydrates for survival. This enzymatic versatility enables the breakdown of plant biomass and has potential for the production of biofuels.^24^ Similarly, *Paenibacillus* species exhibit extensive repertoires of xylan-active CAZyme, allowing the effective degradation of hemicellulose, a primary carbon source in certain environments.^25^ Key genes involved in nitrogen and sulfur cycling were identified, including those responsible for nitrite reduction in nitric oxide and thiosulfate oxidation through the sulfur oxidation (Sox) complex.^26^ Nitrite reduction is a critical step in the denitrification pathway, facilitating the conversion of nitrite to gaseous nitrogen forms and influencing nitrogen availability in ecosystems.^27^ The Sox multienzyme complex, crucial in the oxidation of sulfur compounds, converts thiosulfate to sulfate and plays an essential role in sulfur turnover.^28^ These findings highlight the contributions of the novel bacterial species to nutrient cycling and biogeochemical processes, particularly in extreme and controlled environments such as cleanrooms. In addition, genes associated with SCFA production were identified, such as acetate, propionate, and butyrate, emphasizing their role in maintaining carbon flux under anaerobic or microaerophilic conditions.^29^ The presence of pyruvate metabolism pathways, including its conversion to acetyl-CoA and formate, further underscores the versatility of these species in energy production. ^30^

The study also discovered genes linked to radiation resistance, particularly those that improve DNA repair and transcriptional regulation under stress. These genes were predominantly found in spore-forming isolates, suggesting that sporulation provides added resilience to harsh conditions. For example, *B. pumilus* SAFR-032 demonstrates remarkable resistance to UV radiation and hydrogen peroxide due to efficient responses to DNA repair and oxidative stress.^31^ Similarly, *D. radiodurans*, known for its extreme resistance to radiation, employs sophisticated DNA repair pathways, which these novel species may share.^32^ The presence of biofilm formation genes in both spore-forming and non-spore-forming species indicates a shared strategy for persistence in nutrient-deprived environments.^33^ Biofilm formation not only protects bacteria from environmental stresses, but also improves their survival in oligotrohpic environments such as industrial and clinical setting cleanrooms.^34^ Genomic analysis revealed that alteration of the antibiotic target is the predominant resistance mechanism, along with less common mechanisms such as antibiotic efflux and inactivation. These findings, while computationally derived, require phenotypic validation to confirm resistance expression.^35^ The alteration of the target of antibiotics, a widespread resistance mechanism, involves the modification of the binding sites of antibiotics to reduce efficacy.^36^ Efflux pumps and enzymatic inactivation further contribute to multidrug resistance, underscoring the need for comprehensive phenotypic evaluations to understand the clinical relevance of these resistance genes.^37,38^

BGC analysis identified 12 distinct clusters, with ectoine and *ɛ*-poly-L-lysine being the most prevalent. Ectoine, a compatible solute, is synthesized by specific gene clusters and plays a critical role in osmoregulation and protein stabilization under osmotic stress. Its presence in these novel species suggests improved resilience to fluctuating environmental conditions.^39^ Similarly, *ɛ*-Poly-L-lysine (*ɛ*-PL) is a naturally occurring homopolymer with antimicrobial properties, widely used in food preservation and pharmaceuticals. The biosynthesis of *ɛ*-PL is mediated by specific gene clusters, and its production has been observed in several *Streptomyces* species.^40^ Identification of *ɛ* -PL BGCs in these novel species indicates their potential for *ɛ*-PL production, which could have significant biotechnological applications. Identification of these BGCs with high similarity to known clusters highlights the potential of these novel species to produce bioactive compounds with applications in biotechnology and medicine. More studies are needed to explore the expression and regulation of these clusters to fully understand their functional roles and potential applications. These comprehensive functional annotations provide valuable insight into the ecological roles of these novel bacterial species, their adaptations to extreme environments, and their potential applications in biotechnology. The ability to metabolize diverse substrates, contribute to nutrient cycling, and produce bioactive compounds underscores their significance in both natural ecosystems and controlled environments such as NASA cleanrooms. Moreover, their potential to degrade cleaning reagents used in such environments raises important implications for their management and utilization in industrial processes.

## Conclusion

This study demonstrates that microbial populations in a cleanroom are transient and diverse, and due to the extremely low abundance of these organisms, they are frequently underrepresented in metagenomic analyses. Novel enrichment strategies, targeting the robustness of extremophile microorganisms, when combined with better DNA extraction techniques designed for these hardy microorganisms, would facilitate improved detection. In addition, the appropriate information from the genomes of these rare extremophiles added to reference databases would eventually improve the efficacy of microbial identification and characterization. The genomic characterization of rare microbial species further expands our knowledge of the adaptation of these extremophiles to nutrient-limited environments. Despite considerable challenges, the isolation, identification, and study of extremophiles in cleanroom environments continue to be essential.^41^ This work is vital to industries such as pharmaceuticals, healthcare, and aerospace, where even small traces of problematic microbes can seriously affect human health, compromise product safety, and threaten mission success.

## Materials and methods

### Sample location and sampling

The Jet Propulsion Laboratory (JPL) spacecraft assembly facility (SAF) (∼1,000 m^2^) was regulated to maintain environmental uniformity, which is essential for the integrity of critical operations. This includes keeping the humidity at 30 ± 5% and the temperature within a strict range of 20 ± 4*^◦^*C. Engineers and scientists were trained to follow stringent quality control rules, which is evidence of the high standards maintained in the facility due to the following detailed weekly cleaning procedures.^42^ The SAF was certified according to ISO-6 when the samples were taken, and during the study period of six months from March to August 2016, the number of particles measured reached a maximum of 8,287 particles with a size of 0.5 µm per cubic foot. During the six-month period, several sampling sessions (n = 11) were carried out to collect 98 samples from the cleanrooms of the Mars 2020 mission assembly facility. Sterilized and pre-moistened polyester wipes (9” x 9” size; Texwipe TX1009, NC, USA) were used to collect samples from one-square-meter areas of the cleanroom floors as previously reported.^10,11^ The coordinates and layout of the sampling sites have been cataloged and published previously.^10,11^

### Sample processing

Immediately after sample collection, the wipes were placed in a sterile 500 mL glass bottle containing 200 mL of sterile phosphate-buffered saline (PBS; pH 7.4; Sigma Aldrich, MO, USA) and manually shaken. Shaking vigorously for 30 seconds guaranteed the release of any particles or microorganisms that the wipes had gathered. These samples were then subjected to a concentration process, which reduced their initial volume to 200 ml to a manageable 5 ml. An InnovaPrep concentrating pipette with 0.45 µm hollow fiber polysulfone tips (InnovaPrep Drexel, MO, USA) made for such precision was used to complete this crucial phase. Following concentration, the volume of each sample was precisely determined using a calibrated tare scale and meticulously documented for subsequent analysis. These concentrated samples were used in several experiments that relied on and did not require culture techniques. In particular, spores and cultivable aerobic bacteria were found using cultivation assays to quantify the microbial burden in these samples. Furthermore, to determine the vitality of the microbial components, viability assays were performed, namely the quantitative polymerase chain reaction of adenosine triphosphate (ATP) and propidium monoazide (PMA-qPCR) (data not shown). Details of ATP and qPCR assays have already been published^10^ and are not relevant to this study.

### Cultivation of microbial population

A wide range of culture assays were used to assess microbial diversity at the Spacecraft Assembly Facility (SAF). Following the NASA standard assay (NSA) methodology, aliquots with a precise volume of 425 µl were taken from the 5 mL of concentrated ambient samples and subjected to a standardized heat shock regimen of 80°C for 15 minutes. The purpose of this procedure was to count spore-forming microbes precisely. Simultaneously, non-heat-shocked samples were spread onto tryptic soy agar (TSA). 100 µl was dispensed in quadruplicate to quantify bacterial and spore populations. The TSA agar plates that contain both heat-shocked and not heated samples were incubated at a consistent temperature of 32*^◦^*C. To estimate the bacterial population, colony-forming units (CFU) were carefully counted at 24, 48, and 72 hours and at the end of seven days. Following this extensive analysis, 89 colonies from the non-heat-shocked cohort and 93 colonies from the heat-shocked samples were isolated. Subsequently, these were stored for further research in semi-solid TSA (1/10th concentration).

### Molecular identification

Purification of bacterial isolates was carried out by restreaking the colonies in freshly made TSA medium and the appropriate biomass (∼1 µg wet weight) was harvested after growing at 32*^◦^*C for 2 days. The Mo Bio UltraClean Microbial DNA Isolation Kit (Mo Bio Laboratories, Carlsbad, CA) was used to extract genomic DNA. The 1.5 kb region of the 16S rRNA gene was amplified using established primers 27F (5’-AGA GTT TGA TCC TGG CTC AG-3’) and 1492R (5’-GGT TAC CTT GTT ACG ACT T-3’). The Sanger sequencing settings and amplification conditions were in line with previously published protocols.^3,43^ DNASTAR SeqMan Pro software was used to further process the sequence data, and the reliable SILVA LTP type strain SSU database, version 132, was used for taxonomy identification. Sequence alignment was performed using MUSCLE software, while phylogenetic relationships were determined from a FastTree-built maximum likelihood tree.^44^ A stringent sequence similarity threshold of 98.7% was used in the delimitation of new species,^45^ and all 16S rRNA gene sequences identified have been archived in the GenBank repository under accession numbers MW130960 – MW131089.

Subsequent to 16S rRNA gene-based microbial identification, WGS-based phylogeny was carried out on all 182 pure colonies. After DNA extraction using the ZymoBIOMICS DNA MagBead kit, WGS libraries were performed using Illumina Nextera DNA Flex kits. The sequencing of the whole genome was performed via the NovaSeq 6000 S4 flow cell in paired end, 2 x 150-bp format. FastQC v.0.11.7 and fastp v.0.20.0^46^ were used to exclude low-quality sequences and adapters, which served as quality control for the sequencing reads. The assembly of WGS was attempted using SPAdes v.3.11.1, while assembly statistics were provided including genome size, contigs numbers, and the N50 metric using QUAST v.5.0.2. ^47^

### Taxonomic classification and overall genome-relatedness indices (OGRI) calculation

The completeness and contamination of the assembled genomes were verified using CheckM2 v1.0.2. All genomes showing the completeness of *>* 95% and contamination of *<*5% were grouped together for further analysis. Taxonomic classification was performed by GTDB-Tk v2.4.0, following the classify wf workflow based on ANI-based genome screening.^48^ GTDB-Tk involves the identification of 120 bacterial marker genes using Prodigal^49^ and HMMER,^50^ followed by multiple sequence alignment of conserved regions and maximum-likelihood taxonomic placement. To further validate our classifications, we obtained all validly described representative genomes of all genera identified from the National Center for Biotechnology Information (NCBI) using the command-line tools EDirect^51^ and ‘bit’.^52^ The pairwise average nucleotide identity (ANI) and the average amino acid identity (AAI) between the novel and a representative set of genomes were calculated with FastANI v1.34^53^ and the aai.rb function of the Enveomics toolbox,^54^ respectively. We also predicted digital DNA-DNA hybridization (dDDH) with Formula 2 as described in the Genome-to-Genome Distance Calculator (GGDC) v3.0. ^55,56^

Whenever possible, we used the 16S ribosomal RNA (16S rRNA) gene sequences generated through the amplicon sequencing strategy (MW130960 – MW131089). When not possible, we extracted two additional key conserved gene sequences, 16S rRNA and DNA gyrase subunit B (gyrB), from the annotated genome files generated by Prokka v1.14.5.^57^ Further taxonomic validation was carried out using 16S rRNA sequences that were queried against the NCBI 16S-type strain database using BLAST v2.13.0.^58^ Likewise, gyrB sequences were analyzed using BLAST to verify sequence identity with the closest type strains.

### Screening of novel bacterial species using essential proteins

Five conserved COGs, which encode essential translation and transcription-related proteins, have been used as taxonomic markers for the reconstruction of the phylogenetic relationships of 14 novel species described. COG0081 encodes Ribosomal Protein L1 (RplA), a 234 amino acid subunit of the prokaryotic 50S ribosomal subunit that binds 23S rRNA, contributes to the assembly and stability of the ribosomal protein, and facilitates the release of tRNA from the E site during translation. COG0533 is a 325 amino acid enzyme responsible for the threonylcarbamoylation of adenine at position 37 in tRNA, which is important to maintain the appropriate reading frame and to ensure accurate codon-anticodon pairing during protein synthesis. COG0541 is the Signal Recognition Particle GTPase (Ffh) composed of between 435 and 500 amino acids. The Ffh protein recognizes and binds signal sequences of nascent polypeptides that are directed towards secretion and membrane insertion, and in this regard, the activity of GTPase is essential for proper targeting and translocation into cellular membranes. COG0013 encodes Alanyl-tRNA synthetase (AlaS), a 961 amino acid enzyme that catalyzes the attachment of alanine to its corresponding tRNA (tRNÂAla), a process vital for incorporating alanine into growing polypeptide chains during translation, with proofreading functions to prevent misacylation of tRNA. COG0086 corresponds to the DNA directed RNA Polymerase beta’ subunit (RpoC), approximately 1,400 amino acids in length, forming the catalytic center of the polymerase, involved in DNA binding, unwinding, and RNA transcript elongation.

WGS and metagenomic sequence data were obtained from the National Center for Biotechnology Information (NCBI) and the Integrated Microbial Genomes (IMG) databases for the analyses. NCBI database comprises WGS data from 23,856 isolates and 63,826 metagenome-assembled genomes (MAGs). In addition, the IMG database provides 24,611 MAGs that were not present in NCBI and 80,141 long contigs of metagenomic fragments not publicly available. Traditional methods for defining operational taxonomic units (OTUs) and constructing phylogenies are labor intensive. To address this, CD-HIT was designed as a program to efficiently analyze large data sets.^59^ Processes databases of sequences in FASTA format and results in a set of non-redundant representative sequences together with a cluster file that documents the sequence groups for each representative. CD-HIT uses short-word filtering to calculate sequence similarity, computing the minimum number of identical short substrings shared by two proteins. If the above method does not confirm similarity, it proceeds with a sequence alignment. CD-HIT facilitates improved performance of downstream sequence analyses with minimal loss of information by reducing the database size. In this paper, we applied CD-HIT for the generation of non-redundant protein datasets, thereby facilitating streamlined searches of databases and improving the sensitivity of homology detection. We added five COG genes in FASTA files from 14 novel species to the IMG dataset. CD-HIT clusters at several identities, setting a minimum cut-off threshold coverage at 70%. The CD-HIT cluster at the cutoff point of 98% was examined in relation to the species boundaries of the species as reported in the Genome Taxonomy Database^60^ when the ANI *>*95%. Of all CD-HIT groups based on identity cut-off points 98%, for genes belonging to the COG003 family, the results show closeness to species based on the criteria established from the Genome Taxonomy Database regarding adjusted mutual information analysis.

### Construction of the phylogenetic tree

Genus-level phylogenetic trees were built for the 10 recognized genera, including an appropriate outgroup with GToTree v1.8.6. This tool exploits hidden Markov models (HMM) through HMMER3 v3.4,^50^ to identify sets of single-copy bacterial genes (SCG), then aligned them through MUSCLE v5.1.^61^ For the *Actinomycetota* group, a set of 138 SCGs were used to place seven novel strains within three genera, *Aeromicrobium*, *Cellulomonas*, and *Microbacterium*. In this context, a total of 119 SCGs was used to phylogenetically categorize 18 newly isolated strains from the *Bacillota* phylum into the following seven genera: *Bacillus*, *Brevibacillus*, *Fredinandcohnia*, *Ornithinibacillus*, *Siminovitchia*, *Sporosarcina*, and *Terribacillus*. The protein alignments produced by GToTree were used to generate phylogenetic trees with 1,000 ultrafast bootstrap replicates using the ModelFinder-Plus model ^62^ in IQ-TREE v2.2.0.3.^63^ A bacterial tree of life was generated that included new strains together with 5,031 complete non-anomalous representative bacterial genomes from the NCBI Reference Sequence (RefSeq) database using a universal SCG set.^64^ All trees at the genus level and the bacterial tree of life were visualized and pruned for key nodes with Interactive Tree of Life v6.7.^65^

### Genome annotation and functional profiling

Prokka v1.14.5, a rapid tool for the annotation of prokaryotic genomes was used that makes inferences across a set of several databases about genes and genomic features such as rRNAs and tRNAs, ORFs, among others.^57^ The functional profiling of the genome was performed using cogclassifier v1.0.5 (https://pypi.org/project/cogclassifier/) and DRAM v.0.1.2.^66^ The cogclassifier maps the annotated genes against the Clusters of Orthologous Groups database, and DRAM provides genomes with metabolic profiles. We also decided to determine genes that confer antimicrobial resistance (AMR) based on the robust tool known as the Resistance Gene Identifier v6.0.3, taking advantage of the CARD v3.2.6 database.^67^ RGI employs homology and SNP-based models to predict resistance genes in genomes referenced from the CARD database. Genes with ”Perfect” or ”Strict” matches were included in the AMR profile to ensure a high confidence prediction of antibiotic resistance potential.

### Estimating the abundance and prevalence of novel species in metagenome

Shotgun metagenomic raw read sequences (n = 236 samples) were obtained from NCBI (accession: PRJNA1150505), sampled from the Spacecraft Assembly Facility (SAF) at the Jet Propulsion Laboratory (JPL), California, to assess the abundance and prevalence of novel strains in this controlled cleanroom environment. Of these, 116 samples were treated with propidium monoazide (PMA) to target only viable and intact cells. Fastp v0.22.0^68^ was used for quality control, which eliminated low-quality reads and adapter sequences, thus providing high-quality metagenomic data. Subsequently, we employed MetaCompass v2.0, ^69^ a reference-guided assembly tool, to identify novel strains in the metagenome. MetaCompass performs read alignment to novel genomes using Bowtie2,^70,71^ followed by consensus genome construction using MEGAHIT. Samples without sequence data assigned to the reference genome were not further analyzed. We determined the percentage of reads that corresponded to the novel species and analyzed the breadth of coverage of consensus sequences for each sample, thus informing the distribution of these strains within the cleanroom environment.

### Comparison of secondary metabolite biosynthetic gene clusters

Secondary metabolites (biosynthetic gene clusters, BGCs) in the novel genomes were searched using antiSMASH v7.1.0. ^72^ The functional roles of these gene clusters were predicted using the Minimum Information on a Biosynthetic Gene Cluster (MIBiG) database v3.1.^73^ For comparative analyzes of gene clusters that encode specific secondary metabolites in novel species, known producers, and other isolates from similar environments, we used Clinker available on the CAGECAT web server CAGECAT (https://cagecat.bioinformatics.nl/tools/clinker).^74^ The identity of the precursor proteins was verified by BLAST.

## Data availability

The draft genome sequences of the 182 strains characterized in this study were deposited in NCBI under BioProject PRJNA728748 (HS genomes), PRJNA1136423 (NHS genomes) and PRJNA935338 (*Neobacillus driksii*). The WGS accession numbers are given in Supplemental File S1, and the genome versions described in this document are the first versions. The codes used in this study are available at https://github.com/RamanLab/spore-to-VO.

## Authors’ contribution

K.V. and N.S. managed the Spore to VO (S2VO) microbial strain collection. K.V. and K.R. conceived and designed the study. K.V., S.K.M.S., V.V. generated the draft of the manuscript with contributions from all authors. S.K.M.S., V.V., and P.S. performed the genome assembly, conducted WGS-based phylogenetic placement, comparative genomics, genome annotation, and functional characterization with inputs from K.R., N.K.S., and K.V.. D.W and N.C,K. performed protein-based phylogeny. S.P. and S.K. performed biochemical characterization of the novel species.

## Acknowledgement

The authors thank Ryan Hendrickson, for reviving and purifying the strains, and Zymo Research Corp. for extracting DNA. We thank Chris Mason for the genome sequencing of these strains and Dongying Wu for his valuable assistance in performing the global metagenome analysis. P.S. is a recipient of the Prime Minister’s Research Fellowship (PMRF) from the Ministry of Education, Government of India. S.K.M.S., V.V. acknowledge the Half-Time Teaching Assistantship (HTTA) from the Ministry of Education, Government of India. K.R. acknowledges support from the Wadhwani School of Data Science and AI. All authors read and approved the final manuscript.

## Supporting Information Available

**Supplementary Table S1.** NCBI Accession IDs of 180 strains isolated from Mars 2020 mission Spacecraft Assembly Facility

**Supplementary Table S2.** Bacterial diversity composition, their seasonal, and temporatal distribution of JPL-SAF environmental microorganisms

**Supplementary Table S3.** BioLog characteristics of all novel strains analyzed during this study

**Supplementary Table S4.** Functional COG categories and their gene counts across 25 bacterial strains from this study

**Supplementary Table S5.** Annotated AMR genes and associated attributes of 14 novel bacterial species isolated from Mars 2020 mission Spacecraft Assembly Facility

**Supplementary Table S6.** Biosynthetic gene clusters of novel strains from this study. Each box reflects the percentage of similarity with known BGCs. UN (Unknown) signifies that a BGC was identified, but the similarity couldnt be calculated due to absence of comparable known BGCs. Empty cells represents no BGC was predicted for those genomes

## TOC Graphic

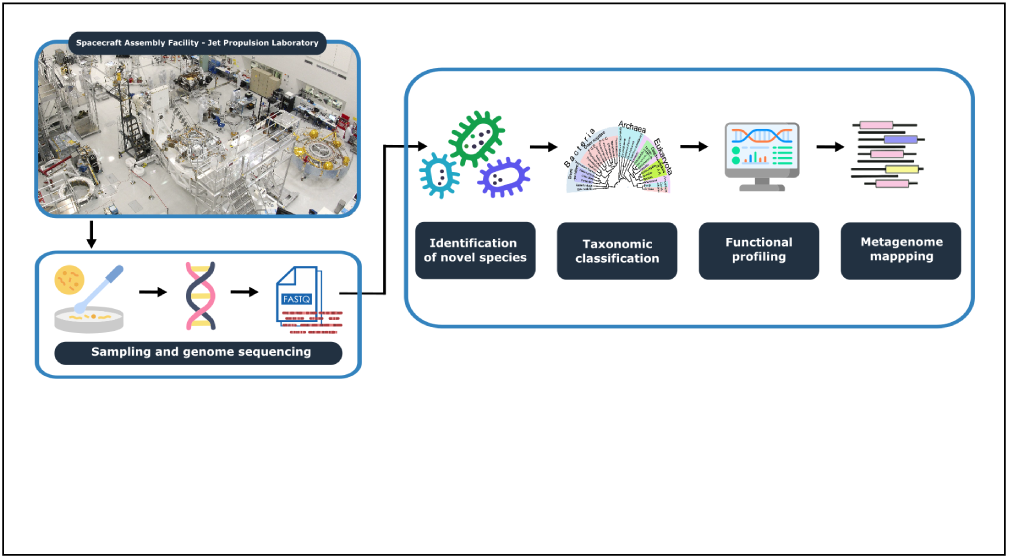

